# Normative assembly rule reveals fairness in microbial communities

**DOI:** 10.1101/2025.11.10.687608

**Authors:** Teemu Kuosmanen, Juhani Rantanen, Dovydas Kičiatovas, Sanna Pausio, Ville-Petri Friman, Teppo Hiltunen, Ville Mustonen

## Abstract

Understanding and predicting how communities assemble is a paramount challenge in ecology. Here we address these questions normatively by comparing the ecological distribution of growth surplus to a game-theoretically fair distribution based on each species’ Shapley value. By analyzing in total 56 distinct community outcomes, we assess how fairly biomass is distributed in microbial communities displaying both competitive and cooperative interactions in different environmental conditions. We find examples of fair communities that closely follow their Shapley value across all environments as well as counterexamples where the true abundances deviate from the species’ objective contribution to community biomass. Our results give unique empirical insights into the distributive function of ecological dynamics and lay down the theoretical foundations of what might become a normative community assembly theory.

## Introduction

Life on Earth thrives in complexity that for now escapes from even proper definition. Organisms interact with each other and their surrounding environment to grow, adapt, and assemble to biological populations, ecological communities, and economic coalitions which in turn self-organize to functioning ecosystems and technological societies with advanced economies (*1*). Such complex systems display various emergent phenomena ranging from cyclicity and chaos (*2, 3*) to robustness and resilience (*4, 5*), and evolution and diversification (*6, 7*). By exploiting available free energy reservoirs, all living organisms produce surplus output that is distributed in the community context by some distinct mechanisms. In nature, the ecological game of growth is largely based on competitive success in utilizing limited shared resources whereas cooperative interactions among social groups, such as seen in human societies, might lead to a different payoff distribution.

By studying competition in its purest form, ecological and evolutionary theory have since their adoption broadly influenced economic thought by shedding light to properties of the ‘state of nature’. While the distribution of income, wealth, and consumption in human societies has been a prominent theme in economic literature (*8, 9*), species abundance distributions (*10, 11*) have similarly been some of the most intensively studied macroecological patterns ever since Preston’s observation that different physical, economical, and ecological ‘wealth distributions’ all seem to take similar form (*12, 13*). But whereas the distributive properties of political and economic systems have also given rise to normative theories addressing questions of distributive justice in terms of fair resource allocation, corresponding ecological theories have remained mainly descriptive, characterizing biodiversity with statistical scaling laws (*14, 15*).

The Shapley value (*16*) is an established solution concept in cooperative game theory, which provides an objective way to distribute the total payoff between the players fairly in the sense that, among all possible payoff distributions, the Shapley distribution is the only one satisfying the mathematical properties of efficiency, symmetry, and linearity (*16*) as well as admitting a unique potential function (*17*). The Shapley value has been extensively applied and extended in the economics game theory literature (*18*) as well as more recently in machine learning as a principled way to quantify the feature importances of different predictors (*19*). In the context of biology, analysis inspired by the Shapley value has found applications for example in the fair proportion index used in constructing phylogenetic trees, to assess importance of genes within a metabolic pathway, or to determine the individual contribution of each taxon to microbiome functional profiles (*20*–*26*).

Despite the extensive exchange of ideas between classical and evolutionary game theory, existing eco-evolutionary theory is predominantly based on applying ideas from non-cooperative game theory focusing on modelling individual strategic behavior. In contrast, cooperative game theory focuses solely on the outputs of different coalitions of players, which are often amenable to direct measurement also in the case of complex systems, where inferring the underlying microscopic interactions is generally very difficult, even for simple minimal models, such as the generalized Lotka-Volterra model commonly employed for studying ecological communities (*27, 28*). Here we demonstrate, perhaps counterintuitively, that concepts of cooperative game theory can be meaningfully applied also in an inherently non-cooperative setting, where competition and exploitation are the norm (*29, 30*).

In this paper, we investigate how biomass as a proxy for cumulative consumption of shared resources is distributed in microbial communities and further pose the question of how fair the resulting ecological outcome is from a normative point of view. By systematically measuring the growth of 14 distinct 4-species communities across all community compositions, we can compute the Shapley value of each species and assess the fairness of the community assembly by comparing these to the actual end-population-shares of the species. We also applied varying antibiotic stress to each community to see whether the observed species abundance distribution and fairness change in more challenging growth conditions, where cooperation can readily emerge via production of public goods, detoxification, or nutrient availability promotion, and thus potentially play a pivotal role in the community outcomes (*31*). By using the established concept of Shapley value, we first empirically quantify the fairness of altogether 56 community outcomes and then develop a normative assembly rule that can also predict unfairly assembled communities if the realized relative abundances in lower level subcommunities are known.

## Results

The Shapley value of player *i* in a game with *n* players forming a grand coalition *N* = {1, …, *n*} is given by (*16*)

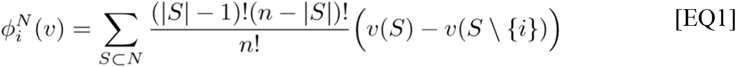

where *v* is a function that gives the output of every *2*^*n*^ possible (sub)coalitions *S* that can be formed from the members of the grand coalition. In the context of ecology, we can interpret *v(S)* as some community function of community *S*, which we assume can be directly measured. Here, we focus on the community yield, denoted by *Y*_*S*_, which is the total biomass at the end of the experiment. Now, to compute the Shapley values, we need not make any assumptions about the ecological dynamics that give rise to the species abundances, but instead simply experimentally measure the observable community yield in all subcommunities (see Fig. 1).

**Fig. 1.**
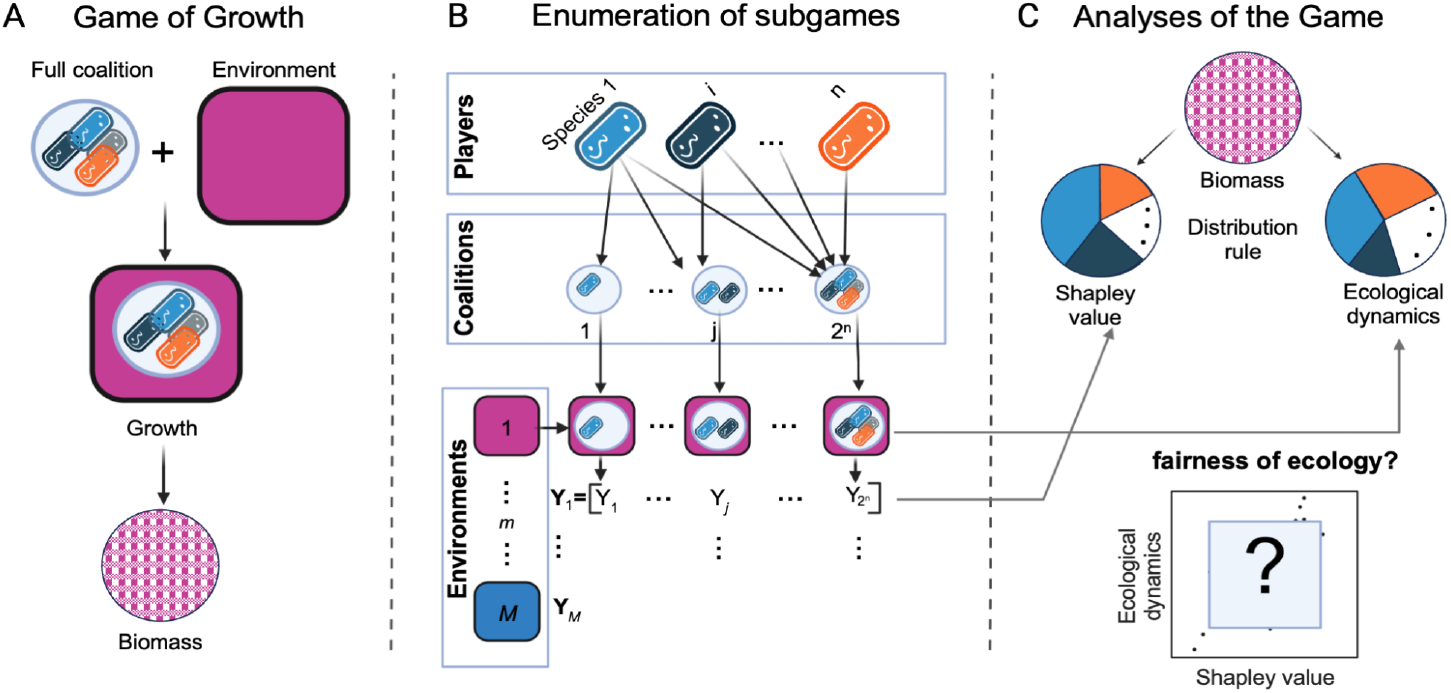
Conceptual overview. (**A)** The observed growth biomass (yield) depends on both the community members, their composition, and the abiotic environment, which together define an instance of *Game of Growth*. (**B)** Here, we measure the growth systematically in all possible community compositions, forming subgames, across all environments, which allows us to quantify the objective contribution of each species by computing its Shapley value. **(C)** By determining the relative proportions of the species in the end by genomic sequencing, we can compare the observed ecological distribution to the game-theoretically fair distribution based on the Shapley value and thus assess the fairness of ecological dynamics as the distribution rule across the communities and environments.

In practice, the Shapley value [EQ1] can be computed more efficiently using the following recursive formula

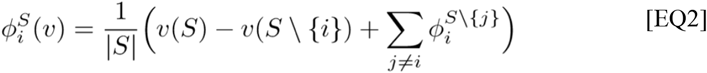

with initial condition 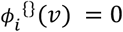 for empty coalitions (for a proof, see (*32*)). This more interpretable formula shows how the Shapley value 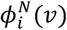 is built bottom-up from the subcoalition Shapley values based on the species’ marginal value on the community function.

We formed 14 distinct communities by randomly selecting four species to each community from a pool of 16 microbial species. We then measured the growth curves and determined the yields from the combinatorially complete set of experiments in four different growth conditions, which allowed us to compute the Shapley values of the community members across the different communities and environments. These results are summarized in Fig. 2.

**Fig. 2.**
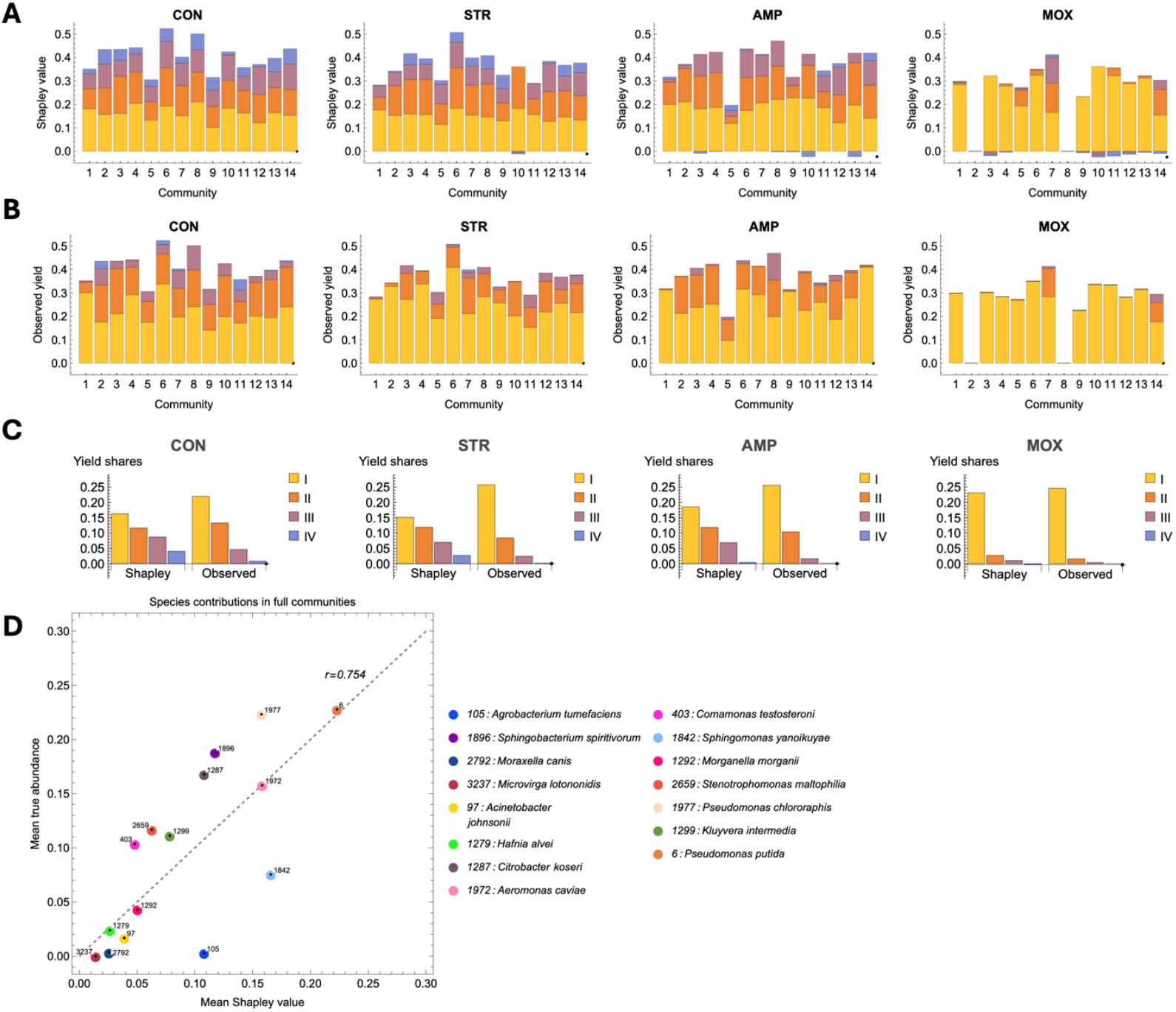
Shapley value analysis of growth across communities and environments. **(A)** The Shapley values of each species in different communities and environments. In each bar, the Shapley values have been independently ranked from largest (I, yellow) to smallest (IV, blue) and the colors refer to the ranking, not species identity. Columns refer to different growth conditions (CON=control condition, AMP=ampicillin, STR=streptomycin, MOX=moxifloxacin). **(B)** The corresponding true abundances of each species ranked in the same way. **(C)** The distribution of yield shares of the ranked four species averaged over all the communities. The observed ecological distribution gives a higher share to the most abundant species compared to their Shapley value. **(D)** The mean Shapley value and true abundance of each species plotted against each other as averaged over all the communities and environments in which the species were present. Species falling below the diagonal receive overall a lower share than what their objective contribution would warrant while the species above the diagonal gain unfairly high abundances.

First, we investigated the yield distribution within each community without focusing on the species identities: by ranking the Shapley values from largest to smallest independently for each community and environment, we gain a first look on how the fair yield shares based on the Shapley value (Fig. 2A) compare to the actual yield distribution (Fig. 2B). Note that by the property of efficiency, the Shapley value distributes the total yield fully among the community members, so we are comparing two different ways to distribute the same surplus output. By averaging these proportions across all communities (Fig. 2C), we noticed that the most prevalent species in each community typically attains higher abundance compared to ranking based on Shapley value, the second ranked species obtains roughly the same share under both distribution rules, and the third and fourth ranked species are left with lower shares than what their Shapley values would warrant. Therefore, the observed yield shares are more concentrated to the most abundant species than the corresponding yield distribution obtained from the Shapley values which would distribute the biomass more equally. However, in the more challenging growth condition of moxifloxacin stress, we noticed that both the Shapley values and the observed yield shares become increasingly concentrated on a single species. In this case, two of the communities produce no observable biomass, and some species even have negative values, meaning that their presence lowered the community output.

We then turned to analyze the roles of the individual species by collecting their Shapley values from all the communities they were present. By the property of linearity, the obtained mean Shapley value thus gives a measure of the species overall deserved share of the games’ output (performance). Although the mean Shapley value of each species is correlated quite strongly (*r*= 0.754) with the true abundances, some species receive on average less than what their objective contribution to the community output is, while others gain a larger share than what could be considered fair (Fig. 2D).

Next, we investigated how closely the observed frequency distribution follows the game-theoretically fair end-distribution *φ*, which can be obtained from the optical density data by normalizing with the sum of the Shapley values, which equals the yield of the grand coalition (see Methods for more details). For example, in the case of only two species, the fair end-distribution is given by

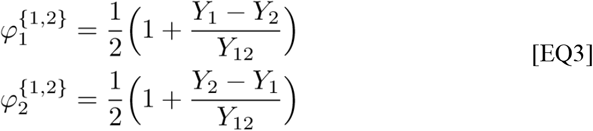

where *Y*_*1*_ and *Y*_*2*_ are the monoculture yields of species 1 and 2 respectively, and *Y*_*12*_ is the biculture yield. This uniquely defines the fair distribution in case of two species for any ecological dynamics, assigning a higher fair proportion for the species with more efficient resource conversion efficiency while the biculture value scales the monoculture difference more strictly in case of antagonism and more loosely in case of synergy allowing for more equal proportions.

In reality, the true end-frequencies after the community growth can deviate from the fair distribution because, in a competitive setting, resource efficiency (monoculture yield) is not the only important factor: for example, shorter lag time, higher resource uptake rate, consumption of secondary metabolites, exploitation of other public goods without reciprocal contribution to their production (cheating), and direct negative interactions, such as bacterial warfare (*29*), can all significantly change the competitive dynamics such that the monoculture yield alone is not predictive enough for success in a community setting. However, species that reach high abundances mainly by exploiting others without contributing themselves cannot grow successfully in harsher environments without the help of the contributing species. Thus, by observing growth systematically across all possible coalitions, we can effectively dissect the causal impact and role of each species on the community output, which defines the fair proportions of the species. By sequencing the community after growth, we can estimate the true end-frequencies, which allows us to compare how ecological reality relates to what would be fair community outcomes.

To quantify the fairness of each community across different environments, we computed the cosine similarity of the fair distribution and the true frequency vector (Fig. 3A). We found communities that systematically followed the fair distribution across all environments; communities that were fair in some conditions, but not others; and communities that deviated significantly from the fair distribution across all conditions. We then quantified both the actual and implied community diversity of the true and fair distributions respectively using the Gini-Simpson index, which can be interpreted as the probability of observing two different species when randomly sampling a pair of individuals from the population, i.e., the probability of interspecific encounter (note that the Gini-Simpson index is inversely related to the Gini index with higher values meaning higher diversity). First, we found that the implied community diversity of the fair distribution could be either high or low, with the latter being the default mode in the presence of moxifloxacin (Fig. 3D). However, in other conditions the fair distribution displayed non-trivial coexistence as showcased in Fig. 3C, which makes the observed fairness of these communities intriguing. We then compared the properties of the fair distribution and the true ecological distribution by contrasting their community diversity against each other (Fig. 3E) and found that the Gini-Simpson index of actual ecological communities was systematically lower or at most as high as that of the fair distribution. This suggests that the ecological reality seems to be more unequal than the fair distribution.

**Fig. 3.**
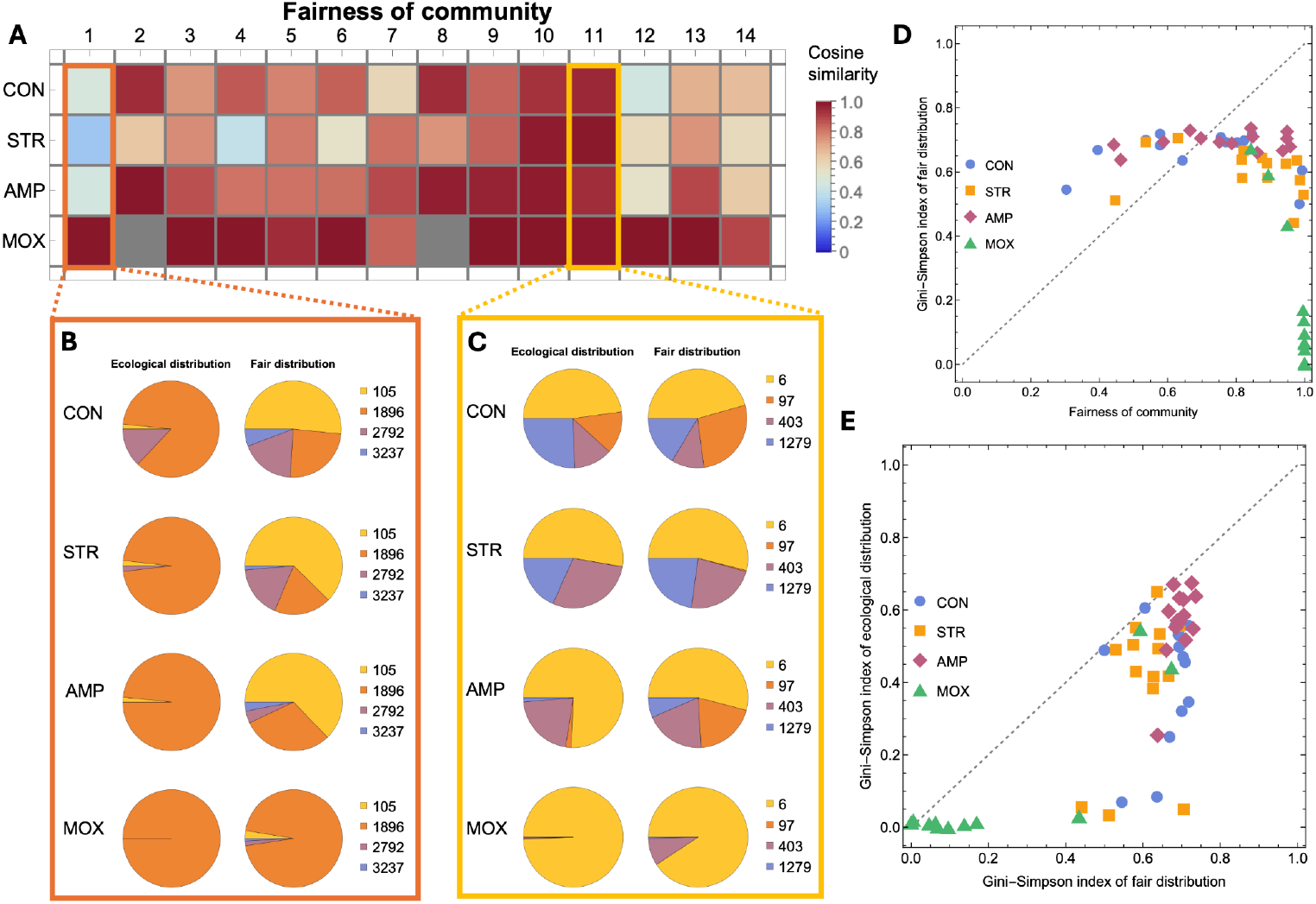
Distributional properties of the ecological versus fair end-distributions. **(A)** We quantified the fairness of each community across all environments by taking the cosine similarity of the true end-frequency vector and the corresponding fair end-frequency vector as given by the normalized Shapley values. Higher cosine similarity means that the community is closer to the fair distribution. Grey color indicates that there was no observed growth in communities 2 and 8 in the MOX-environment. **(B)** Example of an unfair community, which clearly deviates from the fair distribution in all but the MOX-environment. **(C)** Example of a fair community, which closely follows the fair distribution in all growth conditions. **(D)** The fairness of the community is not strongly correlated with community diversity of the fair distribution, and in the stronger selective environment (MOX), the fair end-distribution leads to low diversity. **(E)** The community diversity of the fair distribution plotted against the diversity of the true ecological distribution. All communities fall below the diagonal meaning that the diversity of the fair distribution seems to set an upper bound for the true diversity observed in this experimental system.

A property of the fair distribution is that any discrepancies present with respect to true ecological outcomes already in the biculture competitions get propagated forward via the recursive logic of the Shapley value as shown by [EQ2]. However, if we have access to such additional biologically relevant information besides the yields, can we use this information to predict the full community assembly?

To test this, we sequenced also all the subcommunities and modified the recursive formula [EQ2] to take the estimated true abundances 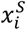 at some subcommunity size *L* as the initial condition for the recursion and then computed the higher-order Shapley values as before. This leads to the following recursive algorithm

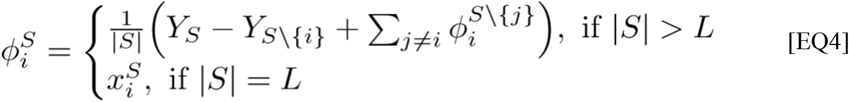

which can be used to predict the species abundances in the full community as 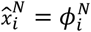. Note that this fair assembly rule contains no free parameters but instead assumes that after the initial unfairness present in the lower subcommunities has been accounted for, the community assembly thereafter follows the rules of the fair assembly.

By incorporating the true biculture and triplet frequencies, we found that the fair assembly rule [EQ4] was able to predict the observed full community outcomes up to very high accuracy (see Fig. 4). We quantified the prediction accuracy using the Euclidean distance between the predicted and observed community composition and found that the mean squared error of the fair distribution with only yields was 0.367 ± 0.034 (SE, N=54). By using the fair assembly rule with the correct pairwise frequencies, this improved to 0.233 ± 0.025 (SE, N=54), and to 0.132 ± 0.015 (SE, N=54) with triplet frequencies. This suggests that the observed assembly process is consistent with the recursive logic of the fair assembly rule.

**Fig. 4.**
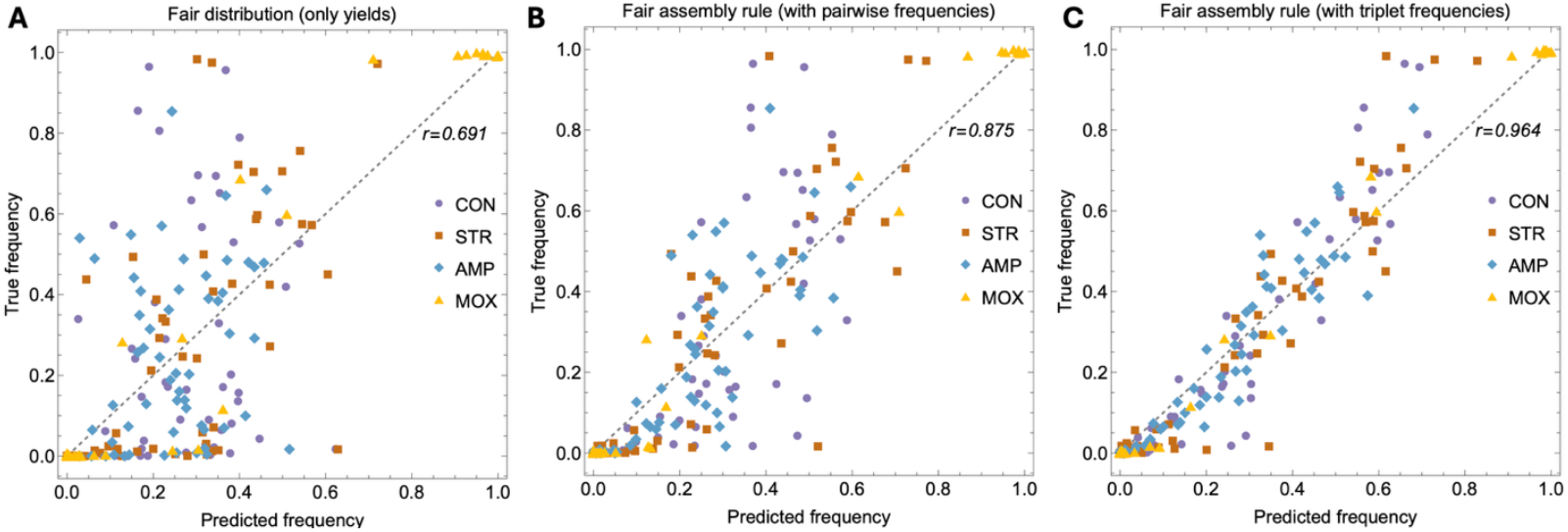
Fair assembly rule predicts community outcomes. The fair assembly rule [EQ4] provides a simple method to predict the full community assembly using the recursive logic of the Shapley value together with information about observed proportions in smaller subcommunities. **(A)** The fair distribution given by the Shapley values captures some aspects of the true assembly process (Pearson’s *r* = 0.691). **(B)** When the fair assembly rule with true rather than fair biculture proportions is used, the full community assembly can be predicted much better (Pearson’s *r* = 0.875). **(C)** When the fair assembly rule with the correct triplet frequencies is used, the predictive performance further improves (Pearson’s *r* = 0.691).

## Discussion

Although much empirical research has been devoted to characterizing biodiversity in terms of species abundance distributions, there is surprisingly little knowledge about the normative status of the ‘state of nature’. Here we asked how fair ecological outcomes are in microbial communities by comparing the observed species abundance distribution to a fair distribution where each species claims only their fair share of the total surplus given by their Shapley value. We found examples of communities where the true ecological distribution followed the fair distribution very closely both in the control conditions and under antibiotic stress, while in other cases some significant species reached unfairly high abundances at the expense of “exploited species” which had a higher objective contribution to the community function than their observed share. One potential mechanism leading to unfairness in our system could be the varying lag time before the onset of growth: we found that species with shorter lag time were generally able to claim their fair share according to their Shapley value while species with a longer lag grew either better or worse after the initial growth of other species (see Fig. S1 in the Supplementary material).

While very unequal payoff distributions can also be fair, as seen in the stronger selective environment (Fig. 3D), we found that the community diversity of the true ecological distribution was systematically lower or at most the same as the corresponding diversity of the fair distribution as measured by the Gini-Simpson index. This tentatively suggests that at least in our microbial model system, the ecological reality seems to be even harsher than that of a purely meritocratic system. However, an important open question remains, whether this applies also in real ecosystems where species have co-evolved together as opposed to the random assemblages used here. Therefore, it may well be plausible that in the absence of ‘unfair’ asymmetries, such as historical priority effects, nature is indeed organized increasingly ‘fairly’ in the Shapley sense. To what extent species sorting and evolutionary mechanisms invoking community state shifts (*33*) can act on fairness remains an intriguing topic for future studies, especially if fairness indeed affects community function. For example, it has been shown that community evenness can facilitate higher robustness in metrics such as invasion resistance and functional redundancy (*34, 35*).

We believe that the introduced framework based on ideas of cooperative game theory and the Shapley value can become broadly useful in analyzing ecological data and developing ecological theory based on the several key strengths highlighted by our work. By construction, the Shapley value compares counterfactual outcomes of the community composition, which allows to make causal inferences of the species contributions as well as their interactions in a model-agnostic way. It can thus provide a general way to make a unique, game-theoretical baseline prediction for community outcomes and as such potentially serve as a useful null model that is driven by experimental data rather than any assumptions about the underlying ecological dynamics. Furthermore, the introduced fair assembly rule [EQ4] could accurately predict complex community outcomes with zero free parameters. This success is particularly interesting, as it remains unclear when microbial assembly follows general rules in contrast to cases where the community member species succeed in “surprising” the experimenters by exhibiting the so-called emergent properties (*36, 37*). Thus, our assembly rule based on fairness opens a new way to identify at what level of hierarchy the possible emergent properties arise. The methodology can also be directly applied to other community functions or properties of interest besides the yield, which was our focus here.

There are, of course, also some limitations to the approach taken here, the most notable being the fact that to compute the Shapley value in practice, one must experimentally measure the yield (or other value of interest) from all the 2^*n*^ possible subcommunities, which grows quickly with the number of present species. However, the technology to make combinatorially complete experiments even for moderately large communities already exists (*38*) and is likely to become easier in the future. Similarly, computer scientists actively develop approximative analysis ideas trying to overcome enumeration challenges related to Shapley values (*24*) which may be transferable to experimental design also in biology. Additionally, if one has access to the realized species distribution at some level, the introduced fair assembly rule can then be used to make predictions for the full community of any size without having to measure all the smaller subcommunities. This makes leave-one-out experiments (see e.g., (*39*)) particularly interesting for testing how these ideas might translate to larger communities.

In conclusion, we have demonstrated how a purely normative concept can be successfully applied to studying a complex system, where a central problem in the typical bottom-up modelling approaches has been the number of unknown parameters that are very difficult to infer even for a minimal model. The suggested approach of using the Shapley value and fairness to understand and predict community assembly is analogous to methods based on optimization principles (*40*) or principle of maximum entropy (*41*), which avoid this problem by deriving the model parameters and predictions from first principles. We anticipate that community assembly theory could be advanced with new types of normative models and identify the evolution of fairness as a particularly interesting area of future research both theoretically related to the evolutionary stability and reachability of fair community compositions as well as empirically in experimental evolution setups.

## Acknowledgments

We would like to acknowledge members of the Bioinformatics and Evolution group for helpful discussions and comments as well as Shane Hogle and Johannes Cairns for their help related to bioinformatics.

## Author contributions

Conceptualization: TK, JR, TH, VM

Formal analysis: TK, VM

Funding acquisition: TH, VM

Investigation: all authors Methodology: TK, JR, DK, VM

Project administration: SP, TH, VM

Resources: TH, VM

Supervision: TH, VM

Visualization: TK, VM

Writing – original draft: TK, JR, VM

Writing – review & editing: all authors

## Materials and Methods

### Sampling Logic and Experimental Setup

In this experiment, we had a pool of 16 bacterial species, from which we randomized a set of 4 species *in silico*. Randomized sets with duplicates of any given strain were discarded, and thus all sets have 4 different species in them. We then formed all possible combinations from these 4 species (n = 15), and this grand coalition is referred to as a block. We formed 25 blocks for this experiment, but sequenced 14 of them, and thus only data from 14 blocks is used in this article.

### Strain Availability and Pre-Culturing

All bacterial strains used in this experiment are from the University of Helsinki Culture Collection (Microbial Domain Biological Resource Centre, HAMBI). For further details on the strains, see supplemental material (Table S1).

Prior to the experiment, we revived the frozen glycerol stocks by axenically cultivating them in 6 ml PPY (proteose-peptone-yeast) medium for 22 hours under experimental conditions. Due to its extended lag phase, we revived HAMBI 3237 in two stages: following an initial 48 h growth period, 500 ul of culture was used to inoculate 6 ml fresh PPY medium for the main pre-cultivation step.

### Growth Assay

Prior to setting up the growth assay, we adjusted the cell density of each culture to 2.4 × 10^7^ cells/ mL by measuring each culture’s OD_600_ and diluting the cultures with M9 minimal medium. The dilution factors were determined based on the relationship between OD_600_ and cell density, established from previously done flow cytometer measurements and CFU (colony-forming unit) counts.

After density adjustment, we assembled the combinations by dispensing 500 µl density-adjusted axenic cultures to the corresponding combination tubes with species number specific volumes of M9. This resulted in the cell density of 6 × 10^6^ cells/ mL per strain in each combination.

For the growth assay, we utilized liquid 100 % R2A (Reasoner’s 2A) medium. In this experiment, we have 4 treatment groups: control, ampicillin, streptomycin and moxifloxacin. The final media antibiotic concentrations were 4.24 µg/ mL for ampicillin, 6.3 µg/ mL for streptomycin, and 0.06 µg/ mL for moxifloxacin. These concentrations are based on median IC50 values across the 16 strains in the bacterial pool the combinations were randomized from. For the control treatment, we added sterile MQ water to dilute the media to have the same amount of nutrients and salts as the antibiotic treatments. The media volume was 140 µl/ well on 96 well cell culture microplates (Corning). We inoculated 10 µl of the assembled bacterial consortia into each well. The growth assays were conducted on Agilent BioTek LogPhase 600 Microbiology Readers at 600 nm, 30 °C for 48 h with shaking on at 800 RPM. The OD_600_ was measured every 10 minutes. After the growth assay, the culture plates were frozen at -20 °C prior to 16S amplicon library preparation.

### Library Preparation

We constructed the 16S amplicon libraries for community composition analysis using an in-house library preparation method described in detail in Cairns et al. (in prep). The method derived from the protocol described in the reference (*42*). Briefly, the V3 region of the 16S gene was amplified with fusion primers which included iTru fusion adapters and an internal sample-specific and combinatorial index. Amplicons were purified with NGS Normalization 96-Well Kit (Norgen Biotek), pooled in sets of 90 reactions, and subsequently amplified with standard TrueSeq primers and indexes. The reaction products were purified with Sera-Mag particles (Cytiva), and equimolarly pooled. The quality of the library was evaluated using Bioanalyzer High sensitivity DNA kit (Agilent) and sequenced on Illumina Miseq at the Finnish Functional Genomics Centre (Turku, Finland), using the MiSeq Reagent Micro Kit v2 (2 × 150 bp reads). DNA template for PCR was obtained by thermal lysis of diluted samples in dH2O (10 min 99,9 °C) and subsequent removal of cellular debris by centrifugation.

### Bioinformatics/ Downstream Sequence Analysis

We demultiplexed the pooled 16S amplicon paired-end sequencing data by: 1) identifying each experimental community sample by the forward and reverse primer sequence combination; 2) trimming paired-end adapter sequences; 3) merging paired-end reads; and 4) filtering merged reads. Scripts to run this pipeline are based on scripts available at https://gitlab.utu.fi/slhogl/hambiDemuxQC.

Identification of samples by primer combinations was done using *cutadapt* (*43*) (version 4.9) software to search sequencing data for primer combinations corresponding to specific experimental communities, as defined during library preparation stage (maximum error rate 0.15). Likewise, *cutadapt* (*43*) (version 4.9) was used to trim paired-end adapter sequences (forward: CCTACGGGAGGCAGCAG; reverse: ATTACCGCGGCTGCTGG; maximum error rate 0.2). Merging of the trimmed paired-end reads was done using *NGmerge* (*44*) (version 0.3) software in default (“stitch”) mode. Filtering of the merged reads was done using *vsearch* (*45*) (version 2.22.1) software (maximum expected error 1; minimum sequence length 115; maximum sequence length 165; “N” characters in sequences are not allowed).

We derived the absolute abundances of the bacterial species within each experimental community using the demultiplexed merged reads with *Rbec* (*46*) (version 1.8.0; as part of R Bioconductor version 3.17) software. Relative abundances obtained by first normalizing species-specific counts by the expected number of 16S gene copies in their genome, then dividing by the total number of counts in the sample. Scripts to run this software are based on the script available at https://gitlab.utu.fi/slhogl/hambiAmplicon.

### Data Analysis

When computing the yields, we subtracted the measured background optical density obtained from the associated control condition. As the Shapley value can become negative for some species, unlike true abundances, we compute the fair end-distribution as *φ*_*i*_ = max(0, *ϕ*_*i*_) / ∑_*k*_ max(0, *ϕ*_*k*_), where the max-operator renormalizes the potentially negative values to give a well-defined fair frequencies that sum up to unity. We quantified community diversity using the Gini-Simpson index defined as 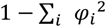.

### Strains in the Bacterial Pool

**Table S1.** HAMBI strains in the pool of 16 bacterial species used in this experiment. Adapted from supplemental material (Table S2) for Cairns et al. 2018. Construction and Characterization of Synthetic Bacterial Community for Experimental Ecology and Evolution. Frontiers in Genetics 9: article 312. DOI: https://doi.org/10.3389/fgene.2018.00312 *Strain 262 was included in the pool, but it was not present in any of the 14 randomized four-species blocks used in the article.

| Strain ID | Species Name | Origin |
| --- | --- | --- |
| 6 | *Pseudomonas putida* | N/A |
| 97 | *Acinetobacter johnsonii* | N/A |
| 105 | *Agrobacterium tumefaciens* | Soil, USA |
| 262* | *Brevundimonas bullata* | Soil |
| 403 | *Comamonas testosteroni* | Soil, Berkeley, USA |
| 1279 | *Hafnia alvei* | N/A |
| 1287 | *Citrobacter koseri* | N/A |
| 1292 | *Morganella morganii* | Human with summer diarrhea, London, UK |
| 1299 | *Kluyvera intermedia* | Surface water, France |
| 1842 | *Sphingobium yanoikuyae* | Clinical specimen |
| 1896 | *Sphingobacterium spiritivorum* | Human, intra-uterine, KS, USA |
| 1972 | *Aeromonas caviae* | Epizootic of young guinea pigs, KY, USA |
| 1977 | *Pseudomonas chlororaphis* | Farm soil, Peoria, IL, USA |
| 2659 | *Stenotrophomonas maltophilia* | Oropharyngeal region of patient with mouth cancer |
| 2792 | *Moraxella canis* | Dog's beard, Finland |
| 3237 | *Microvirga lotononidis* sp. Nov. | Nitrogen-fixing nodule of *Lotononis angolensis*, Chibala, Zambia |

**Fig. S1.**
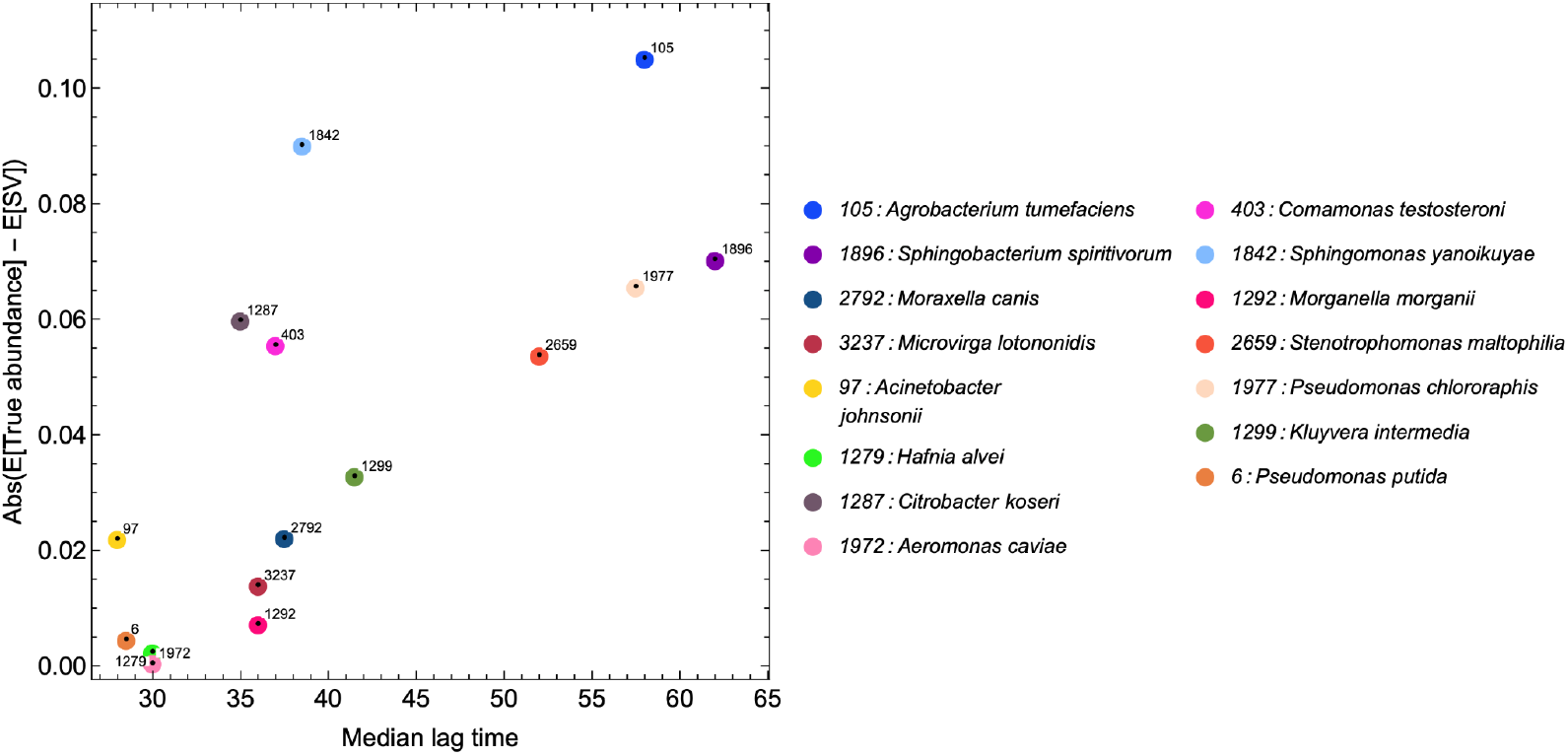
Lag time is correlated with unfairness. Using the optical density measurements we first computed the lag time associated with each growth curve as the time until maximum growth rate. We then plotted the median lag time of each species against the observed unfairness (absolute distance to the diagonal in Fig. 2D). This shows that species with shortest lag time (that is, those that start to grow earlier) are also those which on average obtain their fair share according to their Shapley value, while species with longer lag time reach either unfairly high or low abundances.

